# Spaceflight Affects Neuronal Morphology and Alters Transcellular Degradation of Neuronal Debris in Adult *Caenorhabditis elegans*

**DOI:** 10.1101/2020.11.10.377143

**Authors:** Ricardo Laranjeiro, Girish Harinath, Amelia K. Pollard, Christopher J. Gaffney, Colleen S. Deane, Siva A. Vanapalli, Timothy Etheridge, Nathaniel J. Szewczyk, Monica Driscoll

## Abstract

Extended space travel, such as crewed missions to Mars and beyond, is a goal for both government space agencies and private companies. Research over the past decades, however, has shown that spaceflight poses risks to human health, including negative effects on musculoskeletal, cardiovascular, and immune systems. Details regarding effects on the nervous system have been less well described. The use of animal models holds great potential to identify and dissect conserved mechanisms of neuronal response to spaceflight. Here, we exploited the unique experimental advantages of the nematode *Caenorhabditis elegans* to explore how spaceflight affects adult neurons *in vivo*, at the single-cell level. We found that animals that lived 5 days of their adult life on the International Space Station exhibited considerable dendritic remodeling of the highly branched PVD neuron and modest morphological changes in touch receptor neurons when compared to ground control animals. Our results indicate hyperbranching as a common response of adult neurons to spaceflight. We also found that, in the presence of a neuronal proteotoxic stress, spaceflight promotes a remarkable accumulation of neuronal-derived waste in the surrounding tissues (especially hypodermis), suggesting an impaired transcellular degradation of debris that is released from neurons. Overall, our data reveal that spaceflight can significantly affect adult neuronal morphology and clearance of neuronal trash, highlighting the need to carefully assess the risks of long-duration spaceflight on the nervous system and to develop countermeasures to protect human health during space exploration.

## Introduction

Humankind as long been fascinated by space exploration. Yuri Gagarin first showed in 1961 that humans could survive in space and Neil Armstrong and Edwin ‘Buzz’ Aldrin became the first to walk on the moon in 1969. Now, the International Space Station (ISS), a multinational collaborative station that orbits Earth, has been continuously occupied with crew since November 2000. Into the future, both governmental space agencies and private companies plan to send crewed missions to Mars and beyond. Research to date, however, makes evident that spaceflight poses health risks to the human body, including detrimental effects to musculoskeletal, cardiovascular, and immune systems [1–5]. Particularly striking is the loss of up to 20% of muscle mass in short-duration spaceflights [6, 7], which can be ameliorated by regular physical exercise [8]. Much less is known about the effects of spaceflight on neuronal morphology and function, especially *in vivo* and at the single-neuron level. Given the impracticality of conducting such studies in humans, the use of animal models in which detailed high-resolution neuronal analyses can be performed is invaluable for assessing conserved neuronal responses to spaceflight and developing effective countermeasures to mitigate the consequences of long-duration missions.

The nematode *Caenorhabditis elegans* is a genetic model widely used in neurobiology, aging, and stress research that is amenable for assessing the effects of spaceflight on neuronal biology given its unique advantages: ease of culture in large numbers due to its microscopic size; a transparent body that allows for *in vivo* imaging of multiple tissues, single cells, and even cellular organelles; and a short lifespan (2-3 weeks) that allows for even a short-duration spaceflight mission to correspond to a large portion of the *C. elegans* lifetime. Importantly, the spaceflight-induced muscle atrophy observed in humans is conserved in *C. elegans*, as several studies reported a downregulation of muscular components in space-flown nematodes [9–12]. The aging *C. elegans* musculature also displays the fundamental characteristics of human sarcopenia (the pervasive age-associated loss of muscle mass and strength that contributes to human frailty) [13] and physical exercise in *C. elegans* mimics human exercise by inducing multi-systemic health benefits [14–16]. In addition to muscle changes, space-flown *C. elegans* have altered metabolism and differential expression of longevity-related genes [10–12, 17]. The *C. elegans* nervous system is composed of exactly 302 neurons, each one with a well-described morphology. Thus, transgenic nematodes in which neuronal subpopulations are fluorescently labeled allow for *in vivo* single-neuron assessments that are virtually impossible in the highly complex mammalian brain.

Here we show that adult *C. elegans* that lived 5 days on the ISS exhibit morphological changes in two distinct types of sensory neurons when compared to ground control animals. We identify hyperbranching as a common response of adult neurons to spaceflight. We also studied proteostressed touch neurons with a focus on a formerly unrecognized mechanism by which *C. elegans* neurons can extrude large membrane-surrounded vesicles that contain neuronal waste (e.g. protein aggregates and damaged organelles) [18, 19]. On Earth, the extruded neurotoxic components are, in most cases, efficiently degraded by the surrounding tissues, which in the case studied, is the nematode hypodermis [18, 19]. We find that under spaceflight conditions, proteostressed neurons are associated with a striking accumulation of neuronal debris in the surrounding tissues not apparent in ground controls, indicating a severe dysregulation in the ability to clear neuronal waste in space-flown, middle-aged animals. Our results reveal spaceflight-associated challenges to systemic proteostasis and underscore the need to carefully assess the cellular consequences of long-duration spaceflight as strategies to maintain health are developed.

## Results

### A spaceflight protocol that features adult-only *C. elegans* culture and enables analysis of microgravity impact on a cohort that spans early adult life to middle age

For all experiments presented in this study, we grew *C. elegans* in a liquid culture of S-Basal with freeze-dried *E. coli* OP50 as a food source. Prior to launch, we cultured synchronized populations of young adult nematodes that we loaded into polyethylene (PE) bags with 6.5 mL of liquid culture (~300 animals/bag) on December 2, 2018 (see detailed optimization of culture conditions [20]). Importantly, given our goal of assessing the effects of spaceflight during adult life rather than during animal development, we added 5-fluoro-2′-deoxyuridine (FUdR) to the culture bags to suppress production of progeny and to guarantee that the animals we loaded into the bags would be the same ones we scored upon their return to Earth, as opposed to their progeny. Our samples were loaded into SpaceX CRS-16, the 16^th^ Commercial Resupply Service mission to the ISS, and launched on December 5, 2018 aboard a Falcon 9 rocket. SpaceX CRS-16 docked to the ISS on December 8, 2018 and the *C. elegans* samples, which were kept in cold stowage (8-13°C) once introduced into their flight bags, were transferred to 20°C for five days, beginning on December 9, 2018. After five days, the *C. elegans* culture bags were transferred to −80°C and kept frozen until return to Earth (Fig. 1). For ground control samples, we loaded young adult nematodes into PE bags on December 5, 2018 and exposed *C. elegans* to the same time frame/temperatures as space samples but maintained cultures on Earth. A key aspect of our experimental design is thus that we generated middle-aged nematodes that experienced most of their physiological aging in microgravity (ISS) vs. normal gravity (Earth).

**Figure 1.**
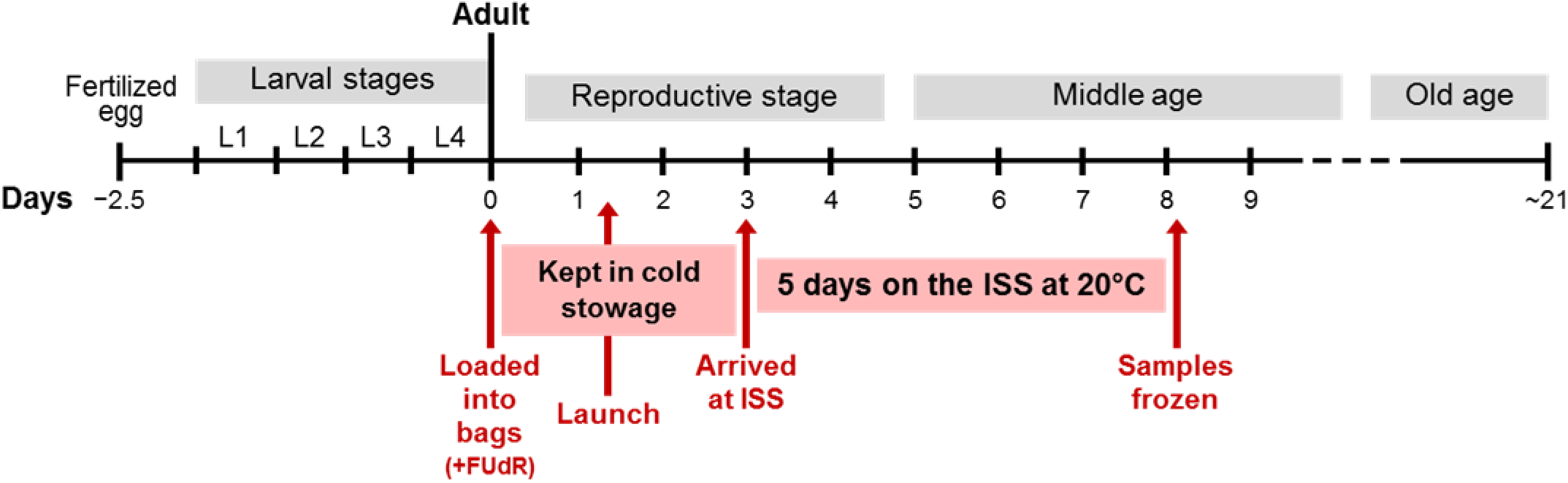
A spaceflight protocol to assess microgravity impact on *C. elegans* adult life. Diagram showing the timeline of *C. elegans* life at 20°C, with approximately 2.5 days of embryonic and larval development and approximately three weeks (21 days) of adulthood. We indicate in red the crucial events in our experimental design by which we obtained middle-aged animals that experienced most of their physiological aging on the International Space Station (ISS). Note that *C. elegans* were kept in cold stowage (8-13°C) for approximately 7 days, relatively low temperatures that significantly delayed progression through adult stages. We estimated the cold stowage period to correspond to approximately 3 days at 20°C. After 5 days on the ISS at 20°C, samples were frozen and returned to Earth for analysis.

### Spaceflight induces morphological remodeling of adult PVD sensory neurons

We started by studying the effects of spaceflight in the morphology of PVD sensory neurons, polymodal nociceptors with spectacular morphologies that sense harsh touch [21], cold temperature [22], and posture [23]. The two PVD neurons (one on the left side, and one on the right side of the body) extend processes that branch to cover most of the body and, together with FLP neurons in the head, possess the most complex dendritic arborization structure of all *C. elegans* neurons (Fig. 2A). The highly branched, yet stereotyped, morphology of PVD dendrites arises sequentially during larval development, with primary (1°) branches forming during the L2 stage, secondary (2°) and tertiary (3°) branches forming during the L3 stage, and quaternary (4°) branches forming during the early L4 stage [24]. By the late L4 stage, PVD patterning is completed and the repetitive structural units resembling candelabras or menorahs become apparent (Fig. 2A).

**Figure 2.**
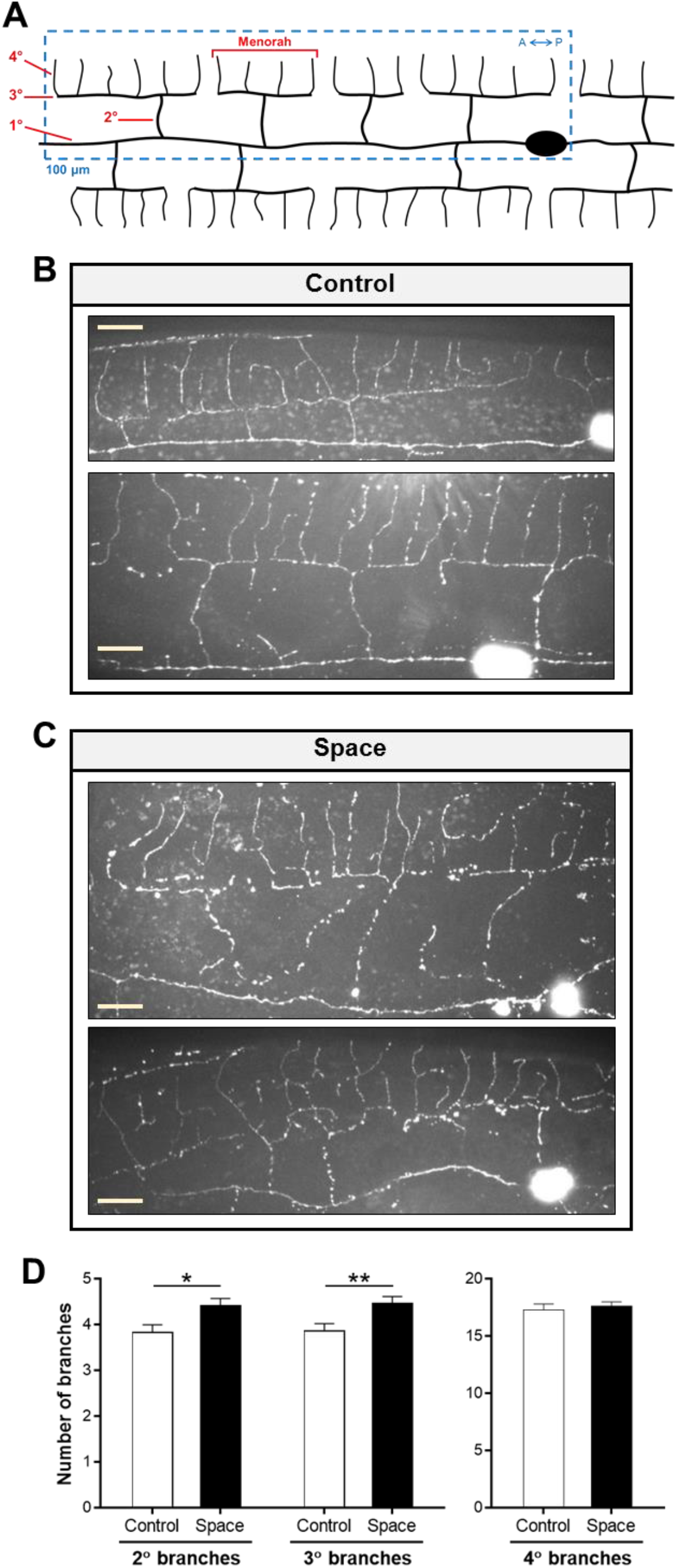
The PVD dendritic tree is affected by spaceflight. (**A**) Diagram showing the morphology of adult PVD sensory neurons. The characteristic PVD branches that form during larval development (1° to 4° branches) to produce the repetitive structural units called menorahs are indicated in red. We performed all quantifications in the 100 μm anterior to the PVD cell body as depicted by the blue dashed box. A, Anterior. P, Posterior. (**B, C**) Representative maximum intensity projection confocal images of ground control (**B**) and spaceflight (**C**) PVD neurons from DES-2::GFP animals. Scale bars, 10 μm. (**D**) Number of 2°, 3°, and 4° branches in ground control and spaceflight PVD neurons. Number of animals used for analysis: *n*_Control_ = 51, *n*_Space_ = 56. We determined statistical significance by unpaired two-tailed Student’s *t* test. \**P* ≤ 0.05, \*\**P* ≤ 0.01.

We used a *C. elegans* strain in which PVD neurons express a translational fusion of transmembrane protein DES-2 and green fluorescent protein (GFP) (DES-2::GFP) [24]. All tiers of the PVD dendrites are well visualized by GFP in this strain and approaches toward quantitative imaging of branching have been published [24–26]. We compared PVD neurons from middle-aged space-flown vs. ground control animals by scoring 1° to 4° branch structures. We found that the space-flown PVD neurons did not degenerate en masse, nor did they exhibit major dendritic gaps relative to ground control PVD neurons (*n* = 51-56 PVD neurons/condition) (Fig. 2B, C). However, when we carefully quantified the number of branches and other menorah-related phenotypes in a 100 μm domain anterior to the PVD soma (Fig. 2A), we registered consistent differences between control and spaceflight animals. More specifically, space-flown animals exhibited increased numbers of 2° and 3° branches, whereas the number of 4° branches remained constant between control and spaceflight animals (Fig. 2D), indicating a plastic morphological restructuring during adult life with outcome that differs consequent to microgravity experience.

On Earth, the PVD dendritic pattern is established during larval stages, but adult neurons still exhibit a limited plasticity of the dendritic trees, with dynamic growth and retraction events occurring during lifetime [26]. Moreover, the PVD dendritic trees show age-dependent hyperbranching, disorganization, and loss of self-avoidance within tree branches [26]. Whether these changes reflect deleterious aspects of aging or adaptive processes as the animal grows remains unclear. With regard to spaceflight samples, our quantification of the number of extra branches derived from 4° branches (5°, 6°, and 7° branches) and the number of ectopic branches (ectopic 2°, 3°, and 4°) found that space-flown animals displayed an increased number of 5° branches (Fig. 3A) and ectopic 3° branches (Fig. 3B) relative to ground control animals. Retrograde branches (4° or higher-order branches that migrate toward the 3° branch by forming a hook) were also present at significantly higher numbers in spaceflight PVD neurons when compared to control counterparts (Fig. 3C). Finally, we observed that spaceflight increased the proportion of disorganized menorahs (menorahs with extra/ectopic branches) (Fig. 3D) and the proportion of self-avoidance defects (no gap in 3° branch between adjacent menorahs) (Fig. 3E).

**Figure 3.**
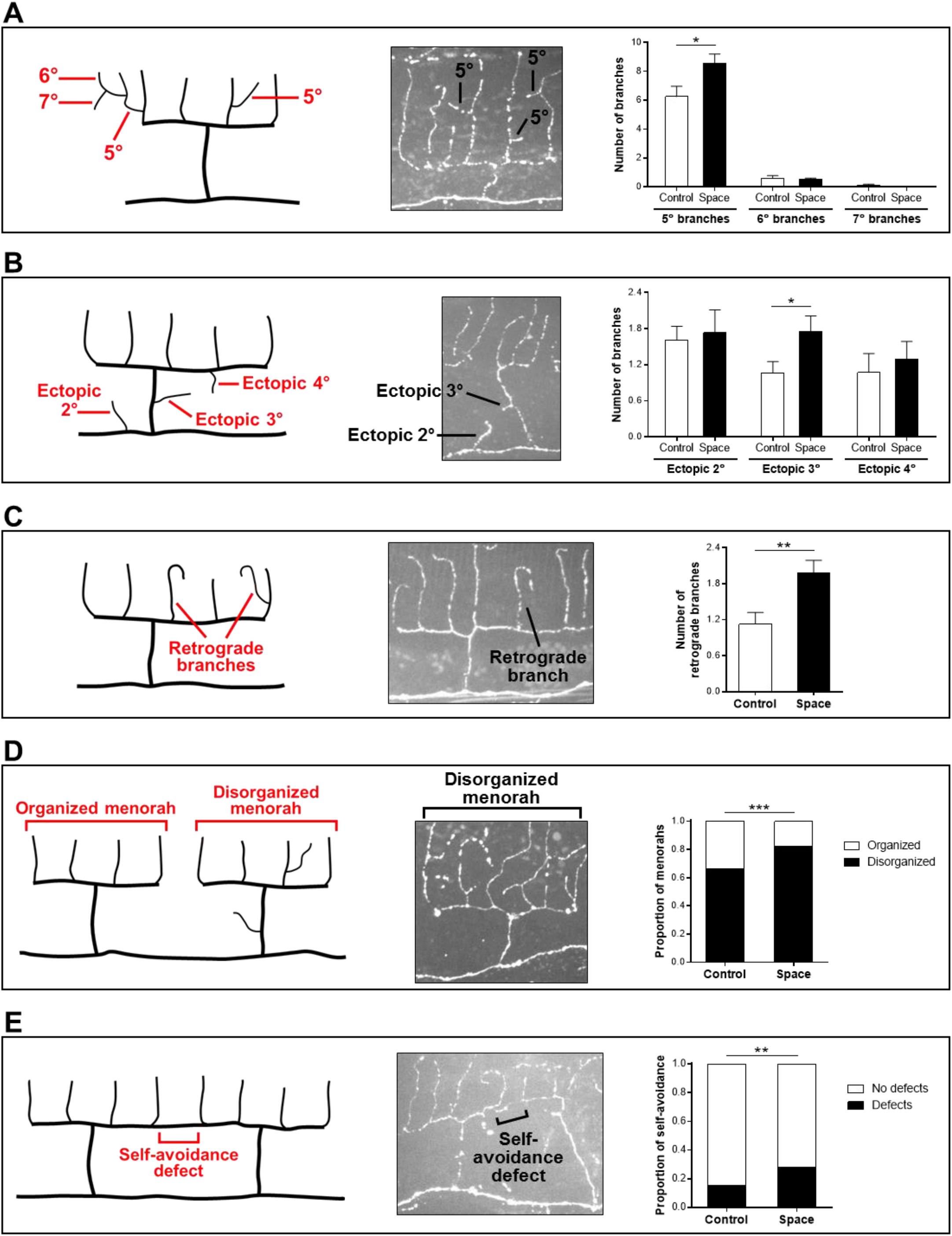
Spaceflight promotes hyperbranching, disorganization, and self-avoidance defects in PVD sensory neurons. (**A-E**) Quantifications in ground control and spaceflight PVD neurons of 5°, 6°, and 7° branches (**A**), ectopic 2°, ectopic 3°, and ectopic 4° branches (**B**), retrograde branches (**C**), disorganized menorahs (**D**), and self-avoidance defects (**E**). We highlight each PVD remodeling phenotype with a diagram (left panel) and a representative confocal image from DES-2::GFP animals (center panel). Number of animals used for analysis: *n*_Control_ = 51, *n*_Space_ = 56. We determined statistical significance by unpaired two-tailed Student’s *t* test (A-C) and by Fisher’s exact test (D, E). \**P* ≤ 0.05, \*\**P* ≤ 0.01, \*\*\**P* ≤ 0.001.

Overall, our data reveal that spaceflight induces significant remodeling of the PVD dendritic tree in adult *C. elegans*. Importantly, the morphological changes we report here for sensory neurons occurred during adult life, indicating that microgravity can influence the morphology of adult neurons.

### Spaceflight promotes modest morphological changes in adult touch receptor neurons

The complex dendritic tree of PVD neurons contrasts with the simple morphology of a single unbranched process adopted by most *C. elegans* neurons [27]. The six touch receptor neurons (AVM, ALML, ALMR, PVM, PLML, and PLMR) mediate the response to gentle touch, with each extending a single major dendritic process anteriorly [28] (Fig. 4A). We sought to investigate the effects of spaceflight on touch receptor neuron morphology by studying a *C. elegans* strain in which touch receptor neurons are specifically labeled with soluble GFP (*P*_*mec-4*_*GFP*).

**Figure 4.**
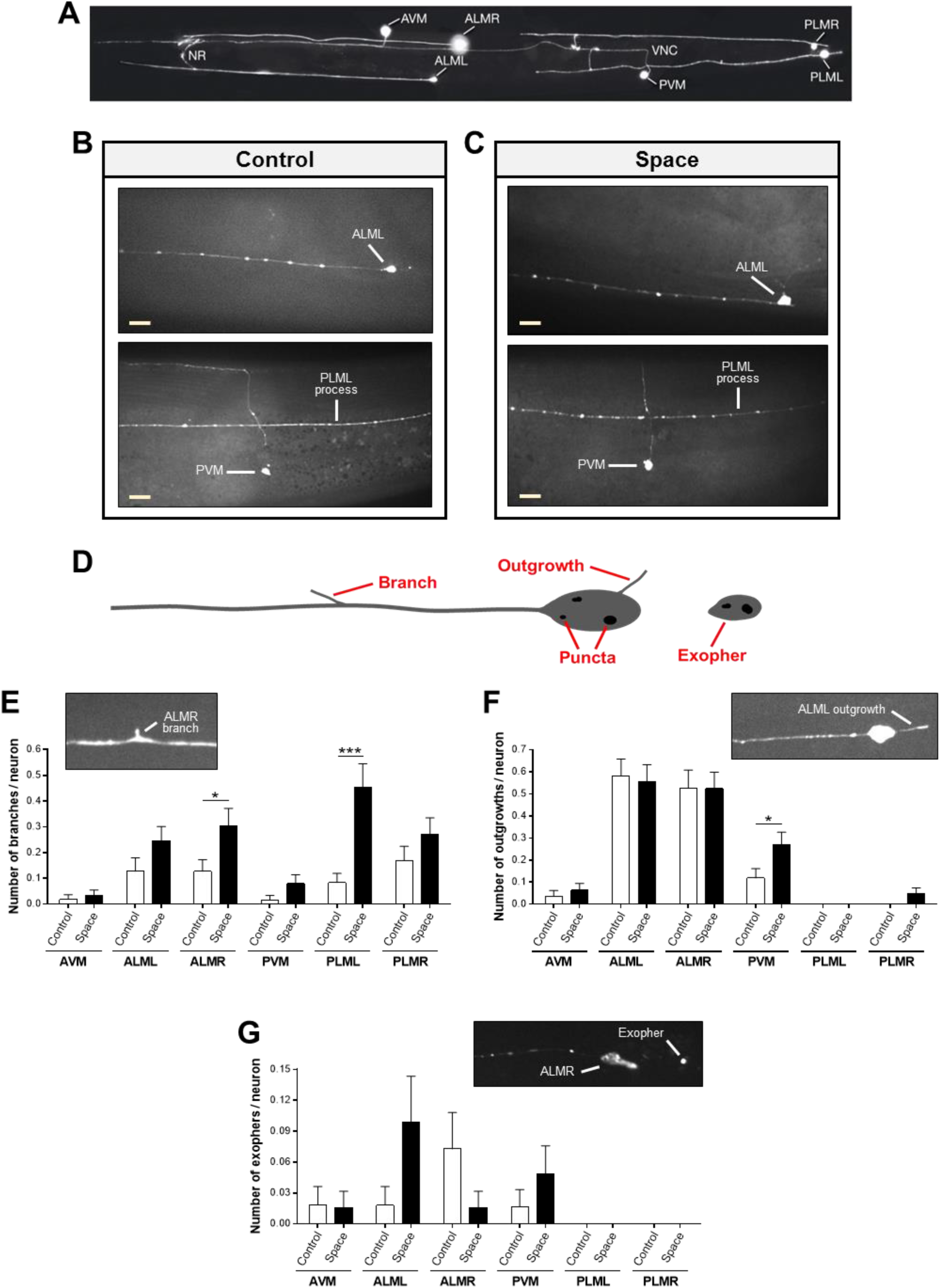
Spaceflight promotes modest morphological changes in adult touch receptor neurons. (**A**) GFP-labeled touch receptor neurons in a young adult *P*_*mec-4*_*GFP* animal. Image adapted from www.wormatlas.org. NR, nerve ring. VNC, ventral nerve cord. (**B, C**) Representative confocal images of ground control (**B**) and spaceflight (**C**) *P*_*mec-4*_*GFP* animals. Scale bars, 10 μm. (**D**) Diagram showing a touch receptor neuron with a branch, an outgrowth, high-intensity fluorescent puncta in the cell body, and an exopher. (**E-G**) Quantifications in ground control and spaceflight *P*_*mec-4*_*GFP* animals of the number of branches (**E**), outgrowths (**F**), and exophers (**G**) in each of the six touch receptor neurons. We highlight each touch receptor neuron phenotype with a representative confocal image from *P*_*mec-4*_*GFP* animals. Number of touch receptor neurons used for analysis: *n*_Control_ = 54-61, *n*_Space_ = 60-64. We determined statistical significance by unpaired two-tailed Student’s *t* test. \**P* ≤ 0.05, \*\*\**P* ≤ 0.001.

We found that middle-aged nematodes that lived five days of adulthood on the ISS exhibited touch receptor neurons with an overall morphology similar to ground control animals, with no obvious degeneration, dendritic gaps, or soma/processes mislocalization (*n* = 54-64 of each touch receptor neuron/condition) (Fig. 4B, C). For a detailed morphological characterization of each neuron, we quantified age-dependent morphological changes previously reported for touch receptor neurons [29], including new branches that extend from the main process (‘branches’) and new processes emerging from the soma (‘outgrowths’) (Fig. 4D). We observed an overall trend in space-flown animals for increased number of branches in all six touch receptor neurons (statistically significant for ALMR and PLML) relative to ground control animals (Fig. 4E), whereas we found the number of outgrowths was generally unchanged by spaceflight (except for PVM) (Fig. 4F). Thus, *C. elegans* adult touch receptor neurons undergo minor structural changes consequent to space travel. Together, our data reveal that spaceflight during adult life does not promote major morphological changes in touch receptor neurons. Still, the hyperbranching phenotype detected in specific adult touch receptor neurons and the morphological changes observed in adult PVD neurons, suggest that modest neuronal restructuring might constitute a general neuronal response to microgravity and/or other stresses associated with spaceflight.

### Spaceflight induces a distinctive response to neuronal mCherry extrusion

Spaceflight includes novel stresses experienced by organisms during lift-off/reentry and extended microgravity periods, and some of these stresses have been suggested to contribute to accelerated aging [30]. Among the many stress response pathways activated, and compromised, by aging are proteostasis pathways such as autophagy and protein degradation processes [31]. Although fluxes through such pathways are difficult to measure in spaceflight samples, we considered some physical readouts thought to reflect proteostasis status.

Expression of GFP in touch receptor neurons can induce high-intensity fluorescent puncta in the soma (Fig. 4D). We found that the absolute numbers of GFP puncta in touch neurons did not differ between middle-aged nematodes on Earth and the ones that spent the corresponding five days on the ISS (Fig. S1A). However, we did observe a trend in space-flown animals for an increased proportion of large GFP puncta (diameter ≥ 0.9 μm) in the soma of touch receptor neurons as compared to ground control animals (Fig. S1B). Although we cannot distinguish whether the touch neuron puncta correspond to aggregates, developing aggresomes, or liquid phase droplets, the differential handling of introduced GFP suggests physiological differences between Earth and spaceflight neurons.

We also reasoned that we might readily evaluate one proteostress response using fluorescence imaging of marked neurons – the extrusion of neuronal garbage. Touch receptor neurons can extrude large membrane-surrounded vesicles called exophers that contain protein aggregates and organelles [19] (Fig. 4D). Exopher production increases under enhanced proteostress as well as under environmental stresses such as elevated superoxide production [18, 19], possibly as a conserved protective mechanism of proteostasis. Under standard conditions, exopher production in GFP-labeled neurons is a relatively rare event [19]. We found that spaceflight did not significantly increase overall exopher detection in touch receptor neurons of *P*_*mec-4*_*GFP* animals (Fig. 4G). We did, however, note an interesting switch in the apparent rates of exopher production in individual touch receptor neurons. On Earth, adult right anterior lateral microtubule neurons (ALMR) reproducibly produce more baseline exophers than other neurons, including the left side counterpart ALML [19], but in spaceflight animals, left side ALML produced more baseline exophers (Fig. 4G). Our observations raise the possibility that maintained gravitational forces may influence left/right asymmetries associated with stresses or their management in particular neurons.

GFP-expressing touch receptor neurons have been studied extensively and are not thought to experience any particular stress as a consequence of the GFP expression [13, 32]. Other transgenically supplied reporters such as mCherry can appear disruptive to normal physiology and are associated with an enhanced level of selective extrusion in exophers [19]. We wondered whether “at risk” neurons, pre-sensitized by the genetic introduction of a noxious mCherry-induced proteostress, might be impacted by spaceflight. Thus, we examined space-flown animals of a strain that over-produces an mCherry reporter well characterized to elevate the production of exophers in touch receptor neurons (*P*_*mec-4*_*mCherry1*) [19]. We examined the neuronal morphology, apparent protein aggregation, and exopher-genesis in the presence of the mCherry stressor. We found that touch receptor neurons of space-flown *P*_*mec-4*_*mCherry1* animals exhibited a generally normal neuronal morphology (*n* = 42-66 of each touch receptor neuron/condition). We observed a trend in spaceflight animals for increased number of mCherry foci in the soma of specific touch receptor neurons (AVM, ALMR, PLML, and statistically significant in PLMR) relative to ground control animals (Fig. S2A), whereas we detected no significant differences in the proportion of large internal mCherry foci (diameter ≥ 0.9 μm) between both conditions (Fig. S2B). We found no significant differences in the absolute number of exophers present in middle-aged control vs. spaceflight touch neurons (Fig. S2C).

We did, however, observe a striking difference in mCherry reporter signal present outside the touch receptor neurons: a remarkably large proportion of *P*_*mec-4*_*mCherry1* animals that spent 5 days of adulthood on the ISS contained numerous mCherry-fluorescent structures throughout the body, a phenotype we never observed in ground control animals (Fig. 5A, B and Fig. S3). While exophers produced by ground control animals are found almost exclusively near touch receptor neuron cell bodies, in limited numbers (one or two exophers produced at the most per neuron), and with characteristic sizes [18, 19], fluorescent structures in space-flown animals ranged from small particles to large rounded structures that could appear as a continuous layer covering multiple regions of the body (Fig. 5B and Fig. S3).

**Figure 5.**
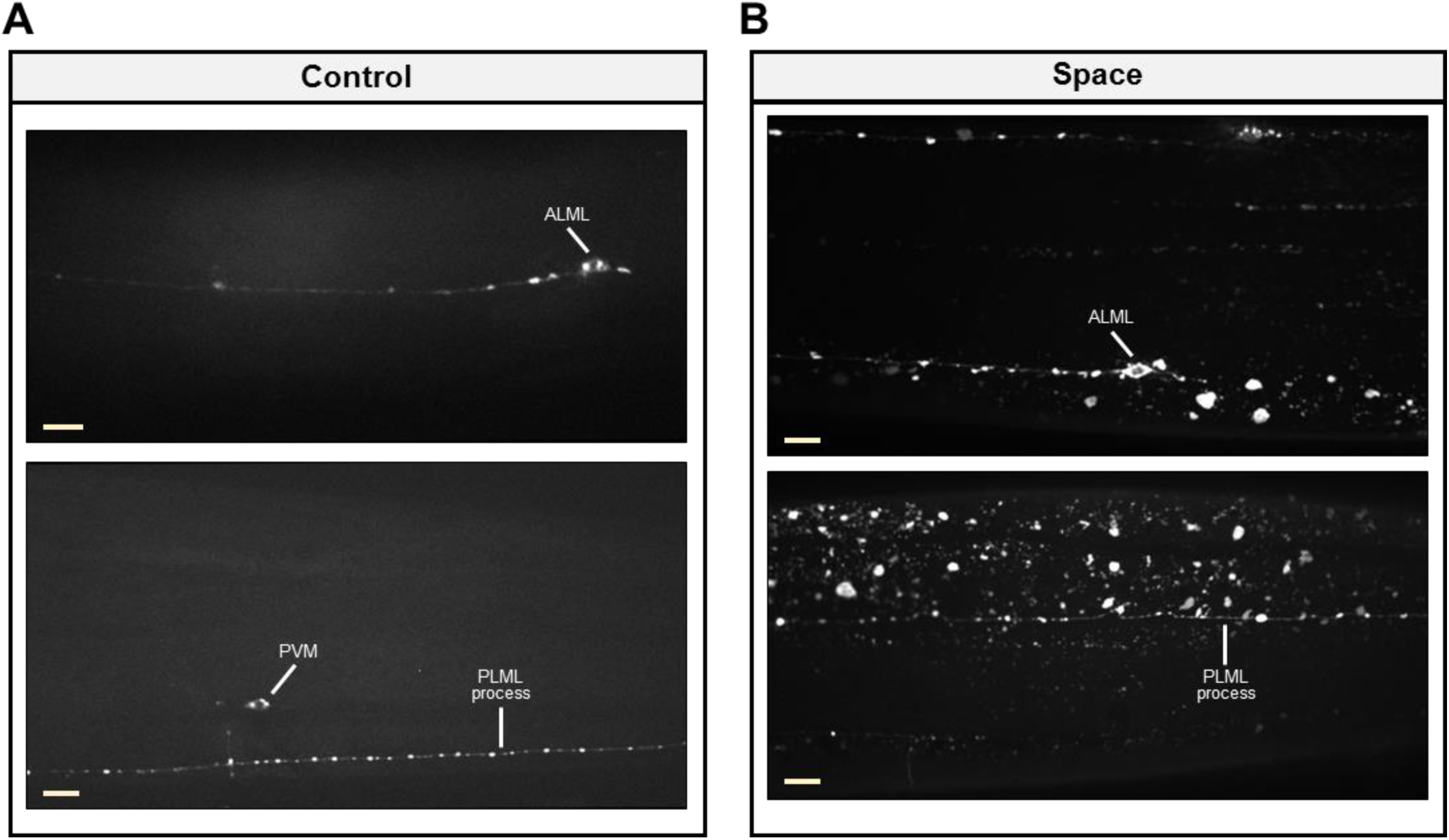
Spaceflight leads to accumulation of neuronal-derived mCherry throughout the body of middle-aged nematodes. (**A, B**) Representative confocal images of ground control (**A**) and spaceflight (**B**) *P*_*mec-4*_*mCherry1* animals. Scale bars, 10 μm.

*C. elegans* touch receptor neurons run through, and are fully surrounded by, hypodermal tissue. When exophers are extruded, they must therefore enter the hypodermis, which attempts digestion of exopher contents via its extensive lysosomal network. mCherry exopher contents that cannot be digested in the surrounding hypodermis are eventually extruded from the hypodermis into the *C. elegans* pseudocoelom. Materials floating through the pseudocoelom can be taken up by scavenger cells called coelomocytes [18, 19]. During the process of transit through the hypodermis, the mCherry exopher contents often become dispersed in the hypodermal lysosomal network, appearing as small fluorescent particles (we refer to this as a ‘Starry Night’ phenotype) [18, 19]. The unexpected distribution of fluorescence that we observed in spaceflight *P*_*mec-4*_*mCherry1* animals appears comprised of two main components: small fluorescent particles resembling the Starry Night phenotype and large rounded fluorescent structures (‘spaceflight vesicles’ or ‘sVesicles’) that we have not previously found in this *C. elegans* strain under standard culture conditions (Fig. 6A). These large rounded structures have the appearance of the enlarged hypodermal lysosomes that can be found in *C. elegans* mutants defective in lysosomal function and assembly [33–35].

**Figure 6.**
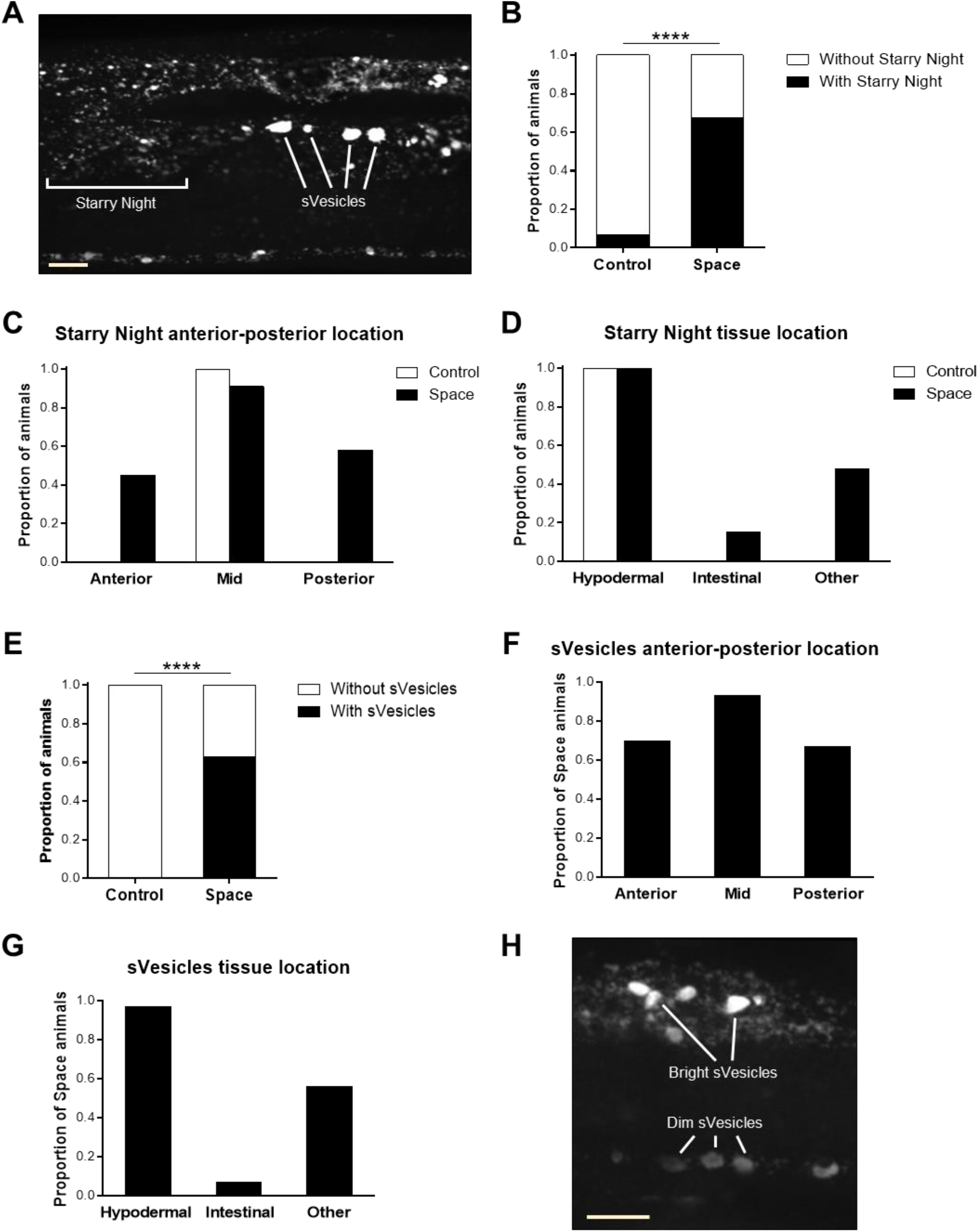
Spaceflight induces a distinctive hypodermal response to neuronal mCherry extrusion. (**A**) Confocal image of a spaceflight *P*_*mec-4*_*mCherry1* animal in which we identified Starry Night and spaceflight vesicles (sVesicles) as two distinct fluorescent components. (**B**) Proportion of ground control and spaceflight *P*_*mec-4*_*mCherry1* animals with/without Starry Night. Number of animals used for analysis: *n*_Control_ = 72, *n*_Space_ = 83. We determined statistical significance by Fisher’s exact test. (**C, D**) Location of Starry Night in ground control and spaceflight *P*_*mec-4*_*mCherry1* animals in the anterior-posterior body axis (**C**) and in different tissues (**D**). Number of animals used for analysis: *n*_Control_ = 5, *n*_Space_ = 33. (**E**) Proportion of ground control and spaceflight *P*_*mec-4*_*mCherry1* animals with/without sVesicles. Number of animals used for analysis: *n*_Control_ = 72, *n*_Space_ = 83. We determined statistical significance by Fisher’s exact test. (**F, G**) Location of sVesicles in spaceflight *P*_*mec-4*_*mCherry1* animals in the anterior-posterior body axis (**F**) and in different tissues (**G**). Number of animals used for analysis: *n*_Space_ = 27. We scored Starry Night and sVesicles as ‘Anterior’ if location was anterior to the AVM neuron, ‘Mid’ if location was in between the AVM and the PVM neurons, and ‘Posterior’ if location was posterior to the PVM neuron. We scored tissue location by imaging different planes on each animal; ‘Hypodermal’ corresponded to the peripheral layer surrounding the body, ‘Intestinal’ corresponded to the cells surrounding the intestinal lumen, and ‘Other’ corresponded to locations in the body not easily identifiable. (**H**) Confocal image of a spaceflight *P*_*mec-4*_*mCherry1* animal showing the variation in fluorescence intensity of sVesicles. Scale bars, 10 μm. \*\*\*\**P* ≤ 0.0001.

We characterized the presence and location of both Starry Night and sVesicles in our samples. We found a striking increase in the proportion of animals with Starry Night in control vs. spaceflight (7% vs. 67%) (Fig. 6B). In ground control animals, we always found Starry Night in the mid-body and in the hypodermis (peripheral layer surrounding the body) (Fig. 6C, D). In space-flown animals, we still found Starry Night predominantly in the mid-body but in approximately half of the cases, the Starry Night dispersed fluorescence expanded into the anterior and/or the posterior region of the body (Fig. 6C). Regarding tissue location, we detected that Starry Night after spaceflight was always present in the hypodermis (Movie S1 and Movie S2) but could also spread into the intestine (15% of animals) or other tissues (48% of animals) (Fig. 6D). We did not observe sVesicles in any of the 72 ground control animals analyzed, whereas 52 out of the 83 space-flown animals (63%) exhibited sVesicles (Fig. 6E). The location distribution of sVesicles in spaceflight animals was similar to Starry Night location, with sVesicles found predominantly in the mid-body but also in the anterior and posterior regions of the body (Fig. 6F), and the primarily hypodermal location (Movie S1 and Movie S2) extended into the intestine (7% of animals) or other tissues (56% of animals) (Fig. 6G). Despite the extensive mCherry fluorescence found outside the touch receptor neurons in spaceflight animals, we rarely observed mCherry fluorescence in coelomocytes in this sample (only 1 out of 56 animals). The similar distribution patterns of Starry Night and sVesicles, together with the fact that 96% of the animals that exhibited sVesicles also had Starry Night, strongly support a link between both phenomena.

We tried to identify the sVesicles found in spaceflight nematodes by performing antibody staining for the lysosomal marker LMP-1 but were unable to get specific staining, possibly due to the paraformaldehyde (PFA) fixation that we had to perform when thawing the frozen nematodes (essential to maintain a strong fluorescence from reporters). Moreover, it appeared that the sVesicles could be readily disrupted as we found that alternative fixation methods (e.g. methanol and acetic acid) led to the disappearance of most mCherry fluorescence outside the touch receptor neurons. The fragility of the sVesicle structures limited our ability to definitively confirm the nature of the hypodermal structures we observed. We did find that the fluorescence intensity of the sVesicles varied greatly, ranging from a bright to a diffuse, dim fluorescence (Fig. 6H), which is consistent with sVesicles being at different stages of lysosomal degradation.

We were fortunate that an unanticipated feature of the flight population enabled us to compare mCherry distribution in some escaper progeny in the culture bag from the ISS. We found that despite our use of FUdR to limit reproduction, some progeny were able to develop up to young adulthood. We could easily identify these animals from older adults by their reduced size. Young adult animals present in the same culture bag as the middle-aged adults that were our major focus exhibited the fluorescent structures at extremely low levels (Fig. S4), suggesting that the dramatic sVesicle profiles in the hypodermis could be age-dependent. We cannot rule out, however, that a variable introduced during development in spaceflight could be responsible for the lack of the large sVesicle pattern in younger animals.

Overall, our data reveal that, in the presence of an internal cell-specific proteotoxic stress (i.e., mCherry overexpression in touch receptor neurons), spaceflight is associated with significant extrusion of the offensive protein in middle-aged touch receptor neurons. Unexpectedly, the extruded material appears to become stuck in the hypodermal lysosomal network, and the lysosomal network adopts a morphology similar to that observed when lysosomes are unable to degrade their substrates. Our data suggest that in microgravity, extruded neuronal garbage is not handled as efficiently by the neighboring cell that collects neuronal debris and attempts degradation.

Future experiments will be necessary to determine the exact mechanism causing mCherry accumulation in large quantities in multiple tissues of *C. elegans* consequent to spaceflight. Still, it is interesting that our results suggest that spaceflight can increase mCherry spread from touch receptor neurons to neighboring tissues (via exophers and/or unknown mechanisms) and that these significant extrusions are not processed in neighboring tissues as occurs on Earth. The abnormal fluorescent structures observed primarily in the hypodermis suggest that the degradative pathways of middle-aged animals may become overwhelmed in response to spaceflight with the resulting inability to quickly clear neuronal trash. Thus, in upcoming spaceflight missions it will be fascinating to specifically track the hypodermal lysosomal network and to determine whether other proteostressors such as aggregation-prone human neurodegenerative disease proteins tau, amyloid beta, huntingtin, or alpha-synuclein, promote similar distinctive outcomes.

## Discussion

Spaceflight has been shown to induce adverse effects on the human body at multiple levels, including in musculoskeletal, cardiovascular, and immune systems [1–5]. However, the effect of spaceflight on neuronal morphology and function, especially *in vivo*, is largely unknown. In this study, we took advantage of the unique characteristics of the nematode *C. elegans* (e.g. ease of culture in large numbers and transparent body allowing for *in vivo* imaging of fluorescently labeled transgenic lines) to address how spaceflight affects the morphology of different adult neurons and the response to a neuronal proteotoxic stress. We show that spaceflight induces extensive remodeling of the PVD dendritic tree in adult *C. elegans* relative to ground control animals, whereas touch receptor neurons exhibit modest morphological changes in response to spaceflight. Importantly, hyperbranching is a consistent response to spaceflight between different adult neuron types. We also report on a striking difference in the ability to respond to a neuronal proteostress (i.e., mCherry overexpression) in spaceflight vs. Earth samples, in which adults that aged on the ISS exhibited accumulated extruded neuronal garbage outside the neurons, distributed in neighboring tissues. Our observations raise the intriguing possibility that spaceflight may impair neuronal proteostasis/clearance pathways, which may hold significant health implications for long-duration spaceflights.

### Spaceflight promotes remodeling of complex adult dendritic trees

We found that spaceflight promoted hyperbranching, disorganization, and loss of self-avoidance in middle-aged PVD neurons. These phenotypes have been shown to increase during aging in *C. elegans* [26], raising the possibility that space-flown animals experienced accelerated aging compared to ground control counterparts. However, adult wild-type animals with disorganized PVD dendritic trees rarely show defects in response to harsh touch [26], suggesting that changes in dendritic architecture are not necessarily a deleterious aspect of aging but rather might be part of normal neuronal maintenance and an adaptive response to intrinsic/extrinsic cues. Furthermore, in other models, most physiological changes induced by spaceflight are reversed upon return to Earth and therefore can be considered physiological adaptations to the spaceflight environment [1, 36, 37]. Thus, we hypothesize that the PVD remodeling we describe here is more likely an adaptive change to spaceflight conditions rather than an overall increase in aging rate. Testing this hypothesis is a challenge for spaceflight experiments as the *C. elegans* lifespan is shorter than typical missions.

A complex crosstalk between muscle, skin (hypodermis), and PVD neurons defines the pattern of higher order branches in the PVD dendritic tree [38–42]. Briefly, the extracellular matrix protein UNC-52/Perlecan forms regular stripes by following the location of the dense bodies of sarcomeres and is essential for the formation of regular hypodermal hemidesmosomes, which are the attachments that mechanically link the hypodermis to the muscle. The intermediate filament MUA-6, which is a component of the hemidesmosome, patterns the transmembrane cell adhesion molecule SAX-7/L1CAM in hypodermal stripes by locally excluding SAX-7 from the regions where MUA-6 is present. Then, SAX-7 physically interacts with the transmembrane receptor DMA-1 in PVD neurons to instruct the growth of 4° branches. Given the well-established and conserved role of spaceflight in muscle degeneration [1, 2, 11, 12], we propose that PVD dendritic morphology in space-flown *C. elegans* may be indirectly affected by changes at the body wall muscle level. In fact, microgravity in *C. elegans* has been shown to downregulate the expression of multiple muscle dense body genes and intermediate filament genes, including *mua-6* [12]. Areas where MUA-6 is absent have abnormally shaped PVD dendrites [39], suggesting that the downregulation of muscle structural genes together with *mua-6* by microgravity could contribute to the changes we observed in space-flown PVD neurons, particularly the hyperbranching, menorah disorganization, and increase in retrograde branches.

Spaceflight has also been suggested to reduce insulin signaling in *C. elegans* [10, 17], whereas the insulin/IGF-1 receptor *daf-2* mutant exhibits an increase in PVD 2° branches [43] and self-avoidance defects [26]. Thus, a putative modulation in insulin signaling in space-flown *C. elegans* might contribute to some of the PVD remodeling phenotypes that we describe in this study.

### Hyperbranching may be a conserved response of adult neurons to spaceflight

We found that space-flown *C. elegans* exhibited touch receptor neurons with modest morphological changes (i.e., hyperbranching) when compared to ground control animals. Interestingly, impaired mitochondrial respiration increases branching in both touch receptor neurons and PVD neurons [43], akin to our observations after spaceflight. Moreover, spaceflight in *C. elegans* has been shown to induce a metabolic shift [10, 11] and decrease the expression of numerous enzymes involved in the tricarboxylic acid (TCA) cycle and of all electron transport chain (ETC) complexes [12], strongly indicating that mitochondrial respiration is reduced during spaceflight. Therefore, changes in mitochondrial function during spaceflight may contribute to the hyperbranching phenotype that we described in different adult neuronal cell types.

The subtle morphological changes induced by spaceflight in touch receptor neurons contrasted with an extensive remodeling of the PVD dendritic tree. This difference in response to spaceflight can be explain by the innate morphology of each neuron (i.e., the highly branched dendritic tree of PVD neurons allows for the detection of multiple morphological abnormalities vs. a single unbranched process of touch receptor neurons), the stimuli sensed by each neuron (i.e., PVD neurons may sense microgravity more directly than touch receptor neurons given their role in proprioception [23]), and/or the fact that PVD morphology is indirectly affected by changes at the muscle level (as discussed previously). Despite the differences between neuron types, our results suggest hyperbranching as a common neuronal adaptive response to spaceflight. This response may in fact be conserved from *C. elegans* to mammals given that Septin 7, a GTP-binding protein critical for dendrite branching [44], has been shown to be significantly upregulated in the brain of mice exposed to the ISS environment for 3 months [45].

### Clearance of neuronal waste is not affected by spaceflight in normal conditions

In addition to neuronal morphological changes, we also assessed how spaceflight modulated the extrusion of neuronal garbage by touch receptor neurons, with a focus on the potential extrusion of large membrane-surrounded vesicles called exophers [19]. We found that GFP-labeled touch receptor neurons, which are not thought to be stressed by transgene expression, exhibited an overall similar number of detectable exophers in middle-aged animals exposed to the ISS vs. Earth environment. It is interesting that for GFP-labeled anterior touch neurons, we measured a difference in the left-right asymmetry in exopher production levels. On Earth, the ALM neuron on the right side of the body (ALMR) consistently produces more exophers than the left side ALM neuron (ALML) [19]. However, in space-flown animals, we observed a reversal in exopher-genesis rate between ALMR and ALML neurons. Although the reason underlying this reversal is unclear, a recent study of reproducible transcriptomic changes in space-flown *C. elegans* [46] identified downregulated genes predicted to be under the control of the transcription factor NSY-7, which is involved in determination of neuronal left/right asymmetry [47]. Together, these results suggest that spaceflight may influence the left/right asymmetries in the *C. elegans* nervous system and potentially affect side-specific neuronal functions.

Exopher production is a relatively rare event under normal conditions (i.e., without a proteotoxic stress) and the majority of exophers are produced on adult days 2 and 3, followed by a reduction in abundance during midlife [18, 19]. In this study, we quantified the number of exophers present in middle-aged animals, a time window several days past the peak in exopher-genesis, and therefore, we cannot definitively draw comparisons to Earth samples regarding the effect of spaceflight on exopher production. We can, however, conclude that spaceflight did not induce any exopher-clearance problems in animals with GFP-labeled touch receptor neurons since we observed no abnormal accumulation of GFP fluorescence in middle-aged *C. elegans*.

### Debris originating in neurons accumulates in surrounding tissues of space-flown animals in the presence of a proteotoxic stress

In the presence of a neuronal proteotoxic stress (i.e., in a line overexpressing mCherry in touch receptor neurons), we observed a surprising accumulation of mCherry fluorescence in the hypodermis (and other tissues) of space-flown animals. Given that mCherry is expressed exclusively in the touch receptor neurons, this fluorescence accumulation in the hypodermis strongly supports that mCherry extruded from touch receptor neurons by exophers and/or other processes is not efficiently cleared in the surrounding hypodermal tissue in space-flown animals.

A major question that arises from our results is whether spaceflight increases the extrusion of neuronal trash in proteotoxic conditions or decreases the degradative/clearance ability of non-neuronal neighboring tissues (or a combination of both). We found a modest increase in the number of mCherry foci in the soma of certain touch receptor neurons exposed to the ISS environment, suggesting that neuronal protein aggregation may be enhanced by spaceflight. This, in turn, could increase the extrusion of neuronal trash (e.g. aggregates) through exophers. We did not identify more exophers in space-flown, mCherry-expressing touch receptor neurons when compared to ground control neurons, but, as discussed above, our quantification occurred in middle-aged animals rather than at the peak of exopher-genesis (adult days 2 and 3). Additionally, given that exopher identification is based on the presence of large fluorescent vesicles nearby the neuron cell body, we likely missed several exophers in space-flown *P*_*mec-4*_*mCherry1* animals due to masking by all the extra fluorescence present. Regardless of a putative increase in neuronal trash extrusion by spaceflight, the remarkable mCherry accumulation throughout the body of over 60% of the space-flown animals indicates to us that the transcellular degradation/management of neuronal trash is severely compromised in the ISS environment.

In mammals, glial cells play a vital role in the central nervous system in the elimination of waste proteins and metabolites produced by neurons [48–50]. These non-neuronal cells maintain homeostasis in the brain and their dysregulation can contribute to the development of neurodegenerative diseases, including Alzheimer’s disease (AD). In fact, microglia and astrocytes play critical roles in the regulation of amyloid-beta (Aβ) clearance and degradation [51]. In the case of *C. elegans*, touch receptor neurons are fully surrounded by hypodermal tissue and therefore, hypodermis assumes an analogous function to mammalian glia regarding the clearance of neuronal waste. Even though we were unable to definitively identify the hypodermal structures in which mCherry fluorescence accumulates in response to spaceflight, it is striking that the large rounded fluorescent structures (sVesicles) resemble enlarged hypodermal lysosomes found in animals defective in lysosomal function and assembly [33–35]. Moreover, the small fluorescent particles (Starry Night) have been previously described as neuron-derived mCherry that becomes dispersed in the hypodermal lysosomal network [18, 19]. Thus, our results suggest that transcellular management of neuronal waste by the hypodermis is defective in space-flown animals, leading to accumulation of impaired degradative organelles such as lysosomes. Our observations also suggest that the inability to efficiently clear neuronal trash is associated with the aging process given the absence of fluorescence accumulation in space-flown young adult animals that were escapers in our study.

On Earth, mCherry extruded from neurons that cannot be digested by the hypodermis is re-released by the hypodermis into the *C. elegans* pseudocoelom and taken up for degradation by scavenger cells called coelomocytes [18, 19]. Interestingly, despite the accumulation of mCherry throughout the body of space-flown animals, we rarely found mCherry fluorescence in coelomocytes, suggesting a defect in hypodermal re-release or an uptake problem in coelomocytes. Alternatively, a distinct debris uptake pathway in other tissues like intestine might be induced in response to spaceflight, so that extruded material is handled differently from neuronal waste on Earth.

Autophagy is a conserved lysosomal-dependent degradative pathway that is critical for cellular homeostasis and has been shown to increase in response to spaceflight or simulated microgravity in multiple cell types, such as muscle [52], bone [53], liver [54], kidney [52], endothelial [55], and cancer [56–58] cells. However, to the best of our knowledge, nothing is known about the effect of spaceflight on autophagy or other degradative pathways in glial cells. Our results may even be consistent with an increase in autophagy markers in the hypodermis of space-flown animals, but the final steps of degradation seem to stall at some point leading to the accumulation of waste-filled vesicles. Future missions should address the impact of spaceflight on glial clearance pathways, particularly in the presence of proteostressors such as neurodegenerative disease proteins tau, amyloid beta, huntingtin, or alpha-synuclein.

### What are the consequences of spaceflight to neuronal function?

In this study, we show that spaceflight can promote significant morphological changes in adult neurons and affect the clearance of neuronal trash from the surrounding tissues. Despite microgravity being the major difference experienced by space-flown animals compared to ground control animals, we cannot exclude other environmental factors (e.g. radiation) as partially responsible for the observed phenotypes. The impact of these changes to neuronal function remains in question; however, our results raise the intriguing possibility that long-duration spaceflights may affect brain function due to neuronal morphological remodeling and deficits in waste degradation. Interestingly, the recent NASA twins study found that the twin astronaut that spent 1 year onboard the ISS exhibited post-flight decline in cognitive performance when compared to his identical brother who remained on Earth [37]. Future missions will need to carefully address the impact of spaceflight on overall brain structure and function to properly assess the risks of long-duration missions planned for the medium future to take humans to Mars and beyond.

## Materials and Methods

### *C. elegans* strains and maintenance

The *C. elegans* strains used in this study were: MF190 *hmIs4[des-2::GFP* + *rol-6(su1006)]* [24], ZB4510 *zdIs5[Pmec-4GFP* + *lin-15(+)] I*, and ZB4065 *bzIs166[Pmec-4mCherry1]* [19]. ZB4510 was generated by outcrossing SK4005 *zdIs5[Pmec-4GFP* + *lin-15(*+*)] I* [59] to N2 five times. Approximately two weeks prior to the launch procedure, we transferred animals from NGM agar plates seeded with live *Escherichia coli* OP50-1 to a liquid culture of S-Basal with freeze-dried *Escherichia coli* OP50 (LabTIE, 1 vial per 250 mL of S-Basal). After this point, we maintained all *C. elegans* strains at 20°C in the liquid culture in 75 cm^2^ cell culture flasks [20].

### Sample preparation for spaceflight

We synchronized *C. elegans* populations by bleaching (20% bleach in 2N NaOH) to obtain eggs or by gravity settling [60] to obtain L1 larvae. When the synchronized animals reached L4/young adult stage, we moved approximately 300 animals per strain to 6-well tissue culture dishes in 1.2 mL of S-Basal with freeze-dried OP50 and 400 μM FUdR. After an overnight incubation at 20°C, we added 5.3 mL of S-Basal with freeze-dried OP50 and 400 μM FUdR to each well and loaded the entire volume (6.5 mL) into PE flight culture bags [20] on December 2, 2018. Culture bags were then placed into experiment cassettes (ECs, Kayser Italia) [20] to provide an additional level of containment and to protect the *C. elegans* cultures from launch and/or in-flight damage. Our samples were then kept in cold stowage (8-13°C) until they reached the ISS. Launch of SpaceX CRS-16 occurred on December 5, 2018 and docking to the ISS occurred on December 8, 2018. On December 9, 2018, our *C. elegans* samples were transferred to 20°C for five days on the ISS. After the five days, samples were transferred to −80°C and kept frozen until return to Earth.

For ground control samples, we performed the same synchronization protocol as for spaceflight samples and we loaded *C. elegans* into the PE flight culture bags on December 5, 2018. We also placed ground control bags into ECs and exposed them to the same time frame/temperatures as spaceflight samples but we maintained cultures always on Earth. Temperature conditions experienced by spaceflight samples were recorded by iButton digital thermometers (iButtonLink) placed inside ECs and allowed us to expose ground control samples to similar conditions. We deposited detailed temperature recordings from spaceflight and control samples in the Harvard Dataverse repository (https://doi.org/10.7910/DVN/ATOJCJ).

### Confocal microscopy

We stored the frozen PE flight culture bags at −80°C until we were ready to start the confocal imaging. We thawed a small fragment of each culture bag in 2% PFA/M9 buffer for approximately 30 min at room temperature. Based on previous testing, we determined that thawing *C. elegans* while simultaneously fixing them in PFA was essential to maintain a strong fluorescence signal in the reporter strains used in this study. After thaw/fixation, we carefully selected, under a stereo microscope, the middle-aged adults present in the sample, which we could easily identify from young animals by their large size. We stored animals in M9 buffer at 4°C until imaging. We performed confocal imaging with an X-Light V2 TP spinning disk unit (CrestOptics) mounted to an Axio Observer.Z1 microscope (ZEISS) using MetaMorph Premier software (Molecular Devices).

### Whole-mount LMP-1 antibody staining

After thawing/fixing *P*_*mec-4*_*mCherry1* animals in 2% PFA/M9 buffer for 30 min at room temperature, we tried different permeabilization methods prior to anti-LMP-1 antibody incubation. We permeabilized animals in 1% Triton X-100/M9 buffer for 45 min, in 1% glacial acetic acid/ethanol for 30 min, or at −80°C in methanol for 1 hour then in acetone for 30 min followed by a serial rehydration at room temperature in 75%, 50%, 25%, and 0% methanol in TBS (100mM NaCl, 50mM Tris-HCl, pH7.5) [61]. We washed samples with 0.1% Tween-20/M9 buffer followed by blocking in 2% BSA/0.1% Tween-20/M9 buffer for 1 hour at room temperature. We incubated samples with anti-LMP-1 antibody (1D4B, Developmental Studies Hybridoma Bank, 1:5 or 1:10 dilution of supernatant) [62] in blocking solution overnight at 4°C. After several washes with 0.1% Tween-20/M9 buffer, we incubated samples with 1:1000 or 1:2000 secondary antibody Goat anti-Mouse, Alexa Fluor 488 (Invitrogen) in blocking solution at room temperature. After washing thoroughly with 0.1% Tween-20/M9 buffer, we imaged samples in the confocal microscope. Despite the different permeabilization methods and antibody concentrations tested, we were unable to observe specific LMP-1 staining in *P*_*mec-4*_*mCherry1* animals.

### Statistical analyses

The data in this study are presented as the mean ± standard error of the mean (SEM). The specific number of data points and the test used for each statistical analysis are presented in the figure legends.

## Acknowledgments

We thank NASA’s Cold Stowage team (payload loading), the Biotechnology Space Support Center (BIOTESC; control of payload operations), and the crew of Expedition 57 (Alexander Gerst, Serena Auñón-Chancellor, and Sergey Prokopyev; payload handling). We thank the UK Space Agency, the European Space Agency (ESA), the Japan Aerospace Exploration Agency (JAXA), and the National Aeronautics and Space Administration (NASA) for their support of the Molecular Muscle Experiment. This work was supported by the BBSRC (Grant Nos. BB/N015894/1) and ESA (designations Molecular Muscle Experiment and ESA-14-ISLRA_Prop-0029). SAV acknowledges support from NASA (Grant No. NNX15AL16G).

---

**Figure S1.**
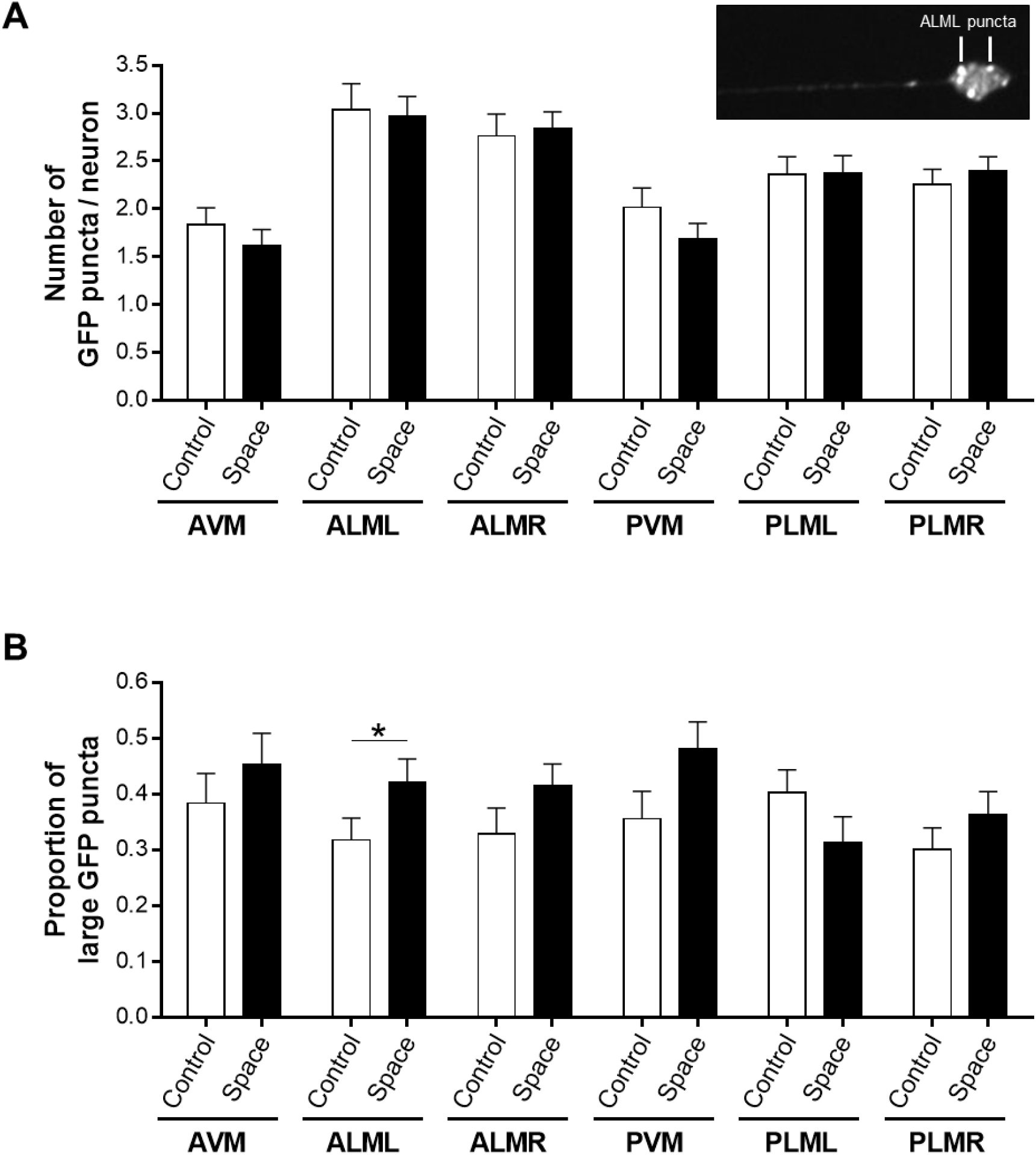
(**A**) Number of GFP puncta per cell body in each of the six touch receptor neurons in ground control and spaceflight *P*_*mec-4*_*GFP* animals. We include a confocal image from a *P*_*mec-4*_*GFP* animal showing GFP puncta in an ALML cell body. Number of touch receptor neurons used for analysis: *n*_Control_ = 54-61, *n*_Space_ = 60-64. We determined statistical significance by unpaired two-tailed Student’s *t* test. (**B**) Proportion of large GFP puncta per cell body in each of the six touch receptor neurons in ground control and spaceflight *P*_*mec-4*_*GFP* animals. We scored GFP puncta as ‘large’ when diameter ≥ 0.9 μm. Number of touch receptor neurons used for analysis: *n*_Control_ = 54-61, *n*_Space_ = 60-64. We determined statistical significance by Fisher’s exact test. \**P* ≤ 0.05.

**Figure S2.**
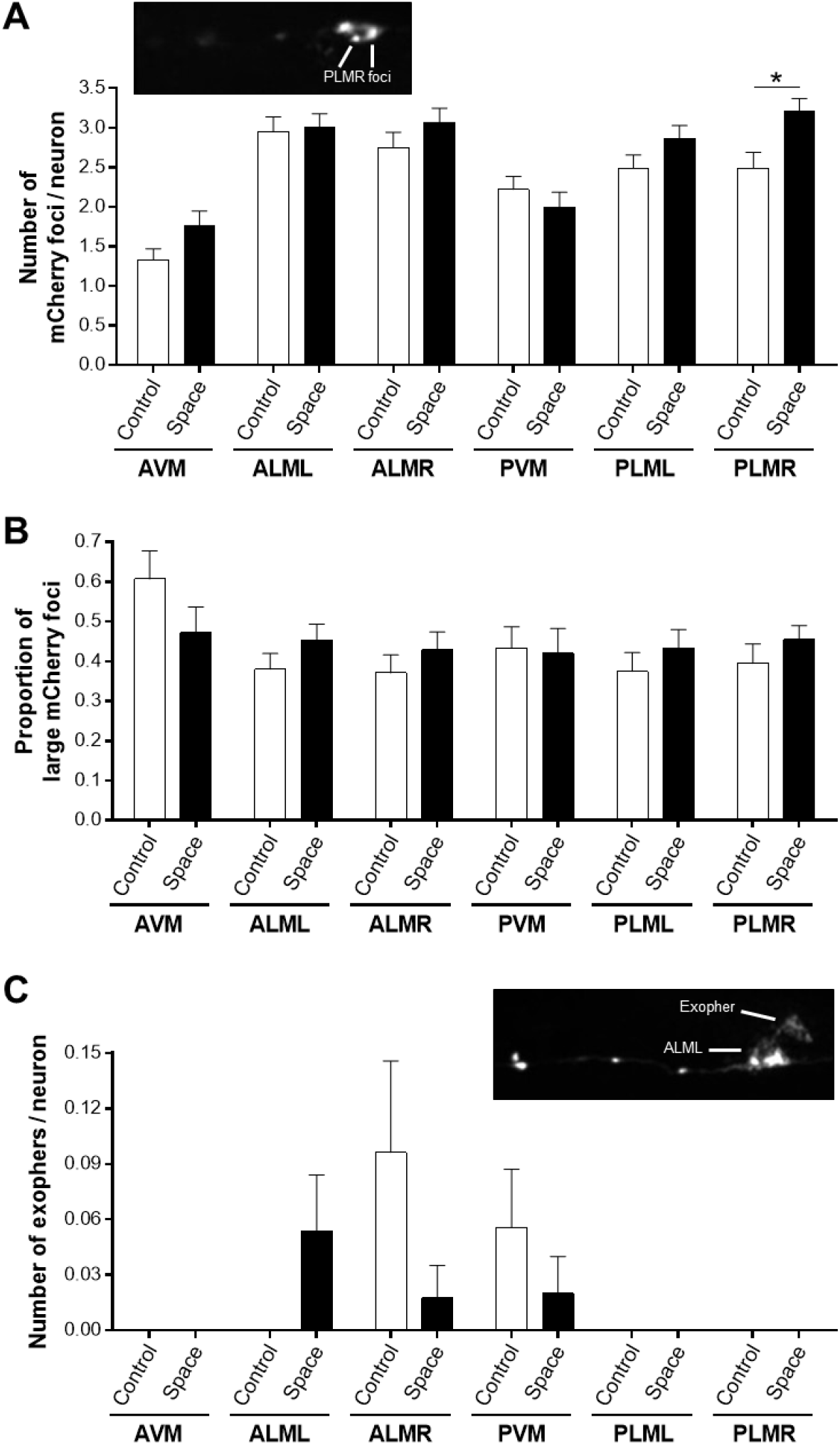
(**A**) Number of mCherry foci per cell body in each of the six touch receptor neurons in ground control and spaceflight *P*_*mec-4*_*mCherry1* animals. We include a confocal image from a *P*_*mec-4*_*mCherry1* animal showing mCherry foci in a PLMR cell body. Number of touch receptor neurons used for analysis: *n*_Control_ = 42-61, *n*_Space_ = 42-66. We determined statistical significance by unpaired two-tailed Student’s *t* test. (**B**) Proportion of large mCherry foci per cell body in each of the six touch receptor neurons in ground control and spaceflight *P*_*mec-4*_*mCherry1* animals. We scored mCherry foci as ‘large’ when diameter ≥ 0.9 μm. Number of touch receptor neurons used for analysis: *n*_Control_ = 42-61, *n*_Space_ = 42-66. We determined statistical significance by Fisher’s exact test. (**C**) Number of exophers per each of the six touch receptor neurons in ground control and spaceflight *P*_*mec-4*_*mCherry1* animals. We include a confocal image from a *P*_*mec-4*_*mCherry1* animal showing an ALML cell body extruding an exopher. \**P* ≤ 0.05.

**Figure S3.**
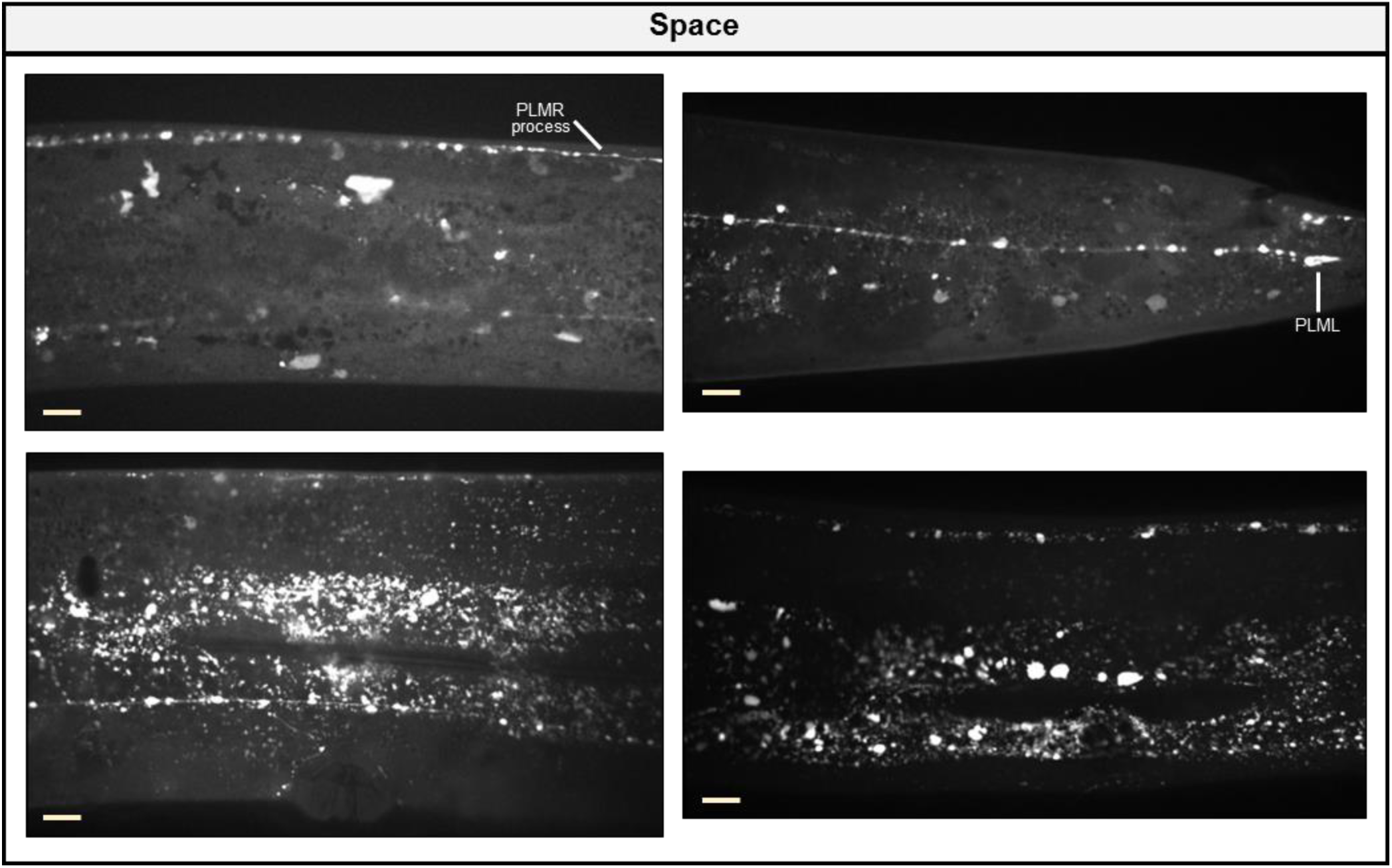
Spaceflight leads to accumulation of neuronal-derived mCherry throughout the body of middle-aged nematodes. Additional representative confocal images of spaceflight *P*_*mec-4*_*mCherry1* animals. Scale bars, 10 μm.

**Figure S4.**
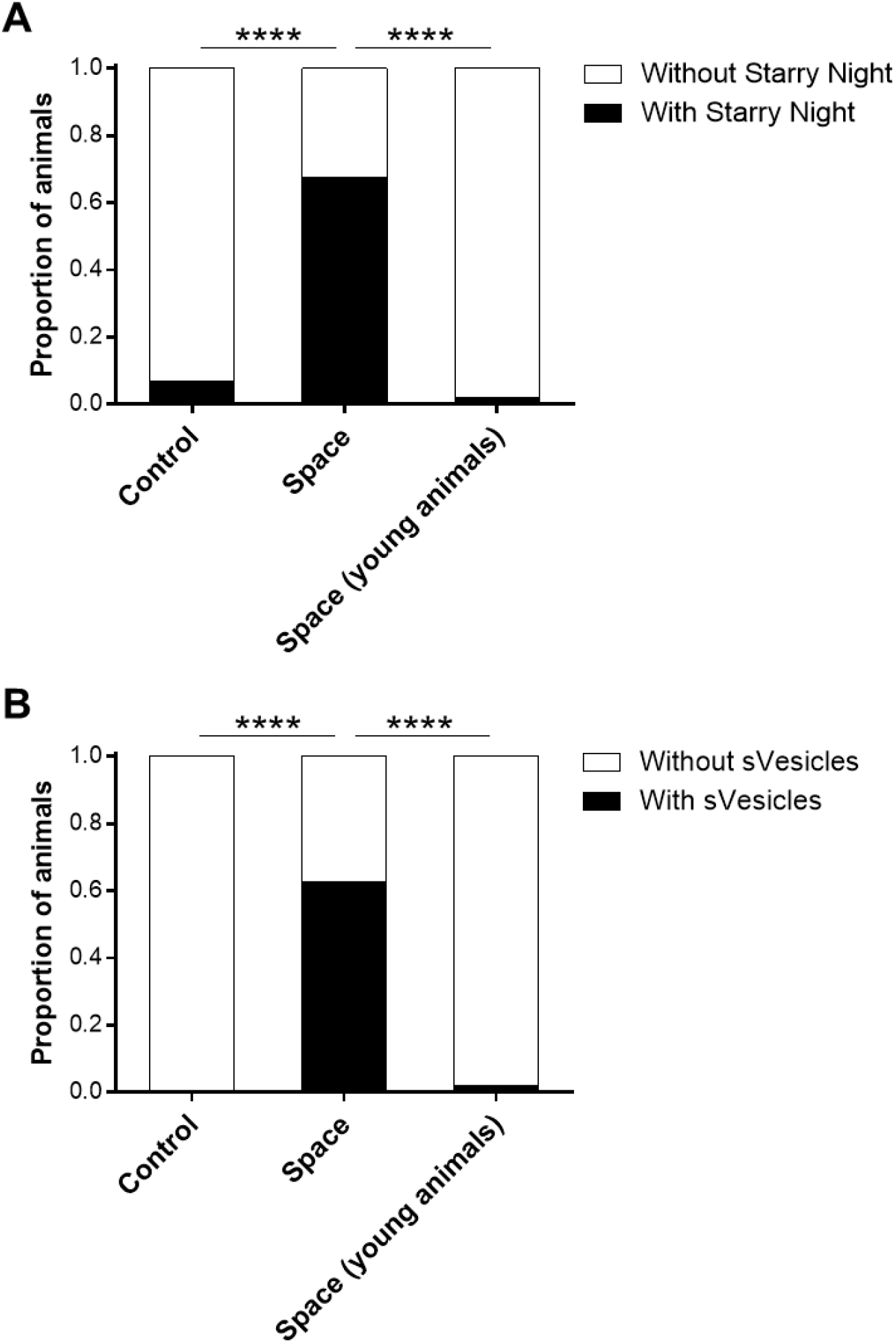
(**A, B**) Proportion of ground control, spaceflight, and young spaceflight *P*_*mec-4*_*mCherry1* animals with/without Starry Night (**A**) and with/without sVesicles (**B**). Note that we obtained young adults from the same spaceflight bag as the middle-aged adults given that some progeny were able to escape FUdR inhibition to develop in the presence of FUdR. We easily identified the spaceflight young adults from the older adults by their reduced size. Number of animals used for analysis: *n*_Control_ = 72, *n*_Space_ = 83, *n*_Space (young animals)_ = 50. We determined statistical significance by Fisher’s exact test. \*\*\*\**P* ≤ 0.0001.

**Movie S1.** 3D reconstruction of the anterior region of a spaceflight *P*_*mec-4*_*mCherry1* animal in which we observed extensive neuronal-derived mCherry (Starry Night and sVesicles) almost exclusively in the hypodermis (peripheral layer surrounding the body). Note that mCherry fluorescence is mostly found on the same focal planes as the touch receptor neurons, which are fully surrounded by hypodermal tissue. AVM, ALML, and ALMR processes are visible in this 3D reconstruction. The animal head is localized toward the left at the start of the movie.

**Movie S2.** 3D reconstruction of the mid-anterior region of a spaceflight *P*_*mec-4*_*mCherry1* animal in which we observed extensive neuronal-derived mCherry (Starry Night and sVesicles) almost exclusively in the hypodermis (peripheral layer surrounding the body). Note that mCherry fluorescence is mostly found on the same focal planes as the touch receptor neurons, which are fully surrounded by hypodermal tissue. The AVM cell body and its process extending downward and then anteriorly are visible in this 3D reconstruction together with the ALMR process (top, same focal plane as AVM) and ALML process (middle, opposite side of AVM and ALMR). The animal head is localized toward the left at the start of the movie.

## REFERENCES

1. Williams, D., et al., Acclimation during space flight: effects on human physiology. CMAJ, 2009. 180(13): p. 1317–23.

2. Fitts, R.H., D.R. Riley, and J.J. Widrick, Functional and structural adaptations of skeletal muscle to microgravity. J Exp Biol, 2001. 204(Pt 18): p. 3201–8.

3. Sibonga, J.D., Spaceflight-induced bone loss: is there an osteoporosis risk? Curr Osteoporos Rep, 2013. 11(2): p. 92–8.

4. Shen, M. and W.H. Frishman, Effects of Spaceflight on Cardiovascular Physiology and Health. Cardiol Rev, 2019. 27(3): p. 122–126.

5. Crucian, B.E., et al., Immune System Dysregulation During Spaceflight: Potential Countermeasures for Deep Space Exploration Missions. Front Immunol, 2018. 9: p. 1437.

6. Edgerton, V.R., et al., Human fiber size and enzymatic properties after 5 and 11 days of spaceflight. J Appl Physiol (1985), 1995. 78(5): p. 1733–9.

7. Adams, G.R., V.J. Caiozzo, and K.M. Baldwin, Skeletal muscle unweighting: spaceflight and ground-based models. J Appl Physiol (1985), 2003. 95(6): p. 2185–201.

8. Loehr, J.A., et al., Physical Training for Long-Duration Spaceflight. Aerosp Med Hum Perform, 2015. 86(12 Suppl): p. A14–A23.

9. Higashibata, A., et al., Decreased expression of myogenic transcription factors and myosin heavy chains in Caenorhabditis elegans muscles developed during spaceflight. J Exp Biol, 2006. 209(Pt 16): p. 3209–18.

10. Selch, F., et al., Genomic response of the nematode Caenorhabditis elegans to spaceflight. Adv Space Res, 2008. 41(5): p. 807–815.

11. Adenle, A.A., B. Johnsen, and N.J. Szewczyk, Review of the results from the International C. elegans first experiment (ICE-FIRST). Adv Space Res, 2009. 44(2): p. 210–216.

12. Higashibata, A., et al., Microgravity elicits reproducible alterations in cytoskeletal and metabolic gene and protein expression in space-flown Caenorhabditis elegans. NPJ Microgravity, 2016. 2: p. 15022.

13. Herndon, L.A., et al., Stochastic and genetic factors influence tissue-specific decline in ageing C. elegans. Nature, 2002. 419(6909): p. 808–14.

14. Laranjeiro, R., et al., Single swim sessions in C. elegans induce key features of mammalian exercise. BMC Biol, 2017. 15(1): p. 30.

15. Hartman, J.H., et al., Swimming Exercise and Transient Food Deprivation in Caenorhabditis elegans Promote Mitochondrial Maintenance and Protect Against Chemical-Induced Mitotoxicity. Sci Rep, 2018. 8(1): p. 8359.

16. Laranjeiro, R., et al., Swim exercise in Caenorhabditis elegans extends neuromuscular and gut healthspan, enhances learning ability, and protects against neurodegeneration. Proc Natl Acad Sci U S A, 2019. 116(47): p. 23829–23839.

17. Honda, Y., et al., Genes down-regulated in spaceflight are involved in the control of longevity in Caenorhabditis elegans. Sci Rep, 2012. 2: p. 487.

18. Arnold, M.L., et al., Quantitative Approaches for Scoring in vivo Neuronal Aggregate and Organelle Extrusion in Large Exopher Vesicles in C. elegans. J Vis Exp, 2020(163).

19. Melentijevic, I., et al., C. elegans neurons jettison protein aggregates and mitochondria under neurotoxic stress. Nature, 2017. 542(7641): p. 367–371.

20. Pollard, A.K., et al., Molecular Muscle Experiment: Hardware and Operational Lessons for Future Astrobiology Space Experiments. Astrobiology, 2020.

21. Way, J.C. and M. Chalfie, The mec-3 gene of Caenorhabditis elegans requires its own product for maintained expression and is expressed in three neuronal cell types. Genes Dev, 1989. 3(12A): p. 1823–33.

22. Chatzigeorgiou, M., et al., Specific roles for DEG/ENaC and TRP channels in touch and thermosensation in C. elegans nociceptors. Nat Neurosci, 2010. 13(7): p. 861–8.

23. Albeg, A., et al., C. elegans multi-dendritic sensory neurons: morphology and function. Mol Cell Neurosci, 2011. 46(1): p. 308–17.

24. Oren-Suissa, M., et al., The fusogen EFF-1 controls sculpting of mechanosensory dendrites. Science, 2010. 328(5983): p. 1285–8.

25. Oren-Suissa, M., et al., Extrinsic Repair of Injured Dendrites as a Paradigm for Regeneration by Fusion in Caenorhabditis elegans. Genetics, 2017. 206(1): p. 215–230.

26. Kravtsov, V., M. Oren-Suissa, and B. Podbilewicz, The fusogen AFF-1 can rejuvenate the regenerative potential of adult dendritic trees by self-fusion. Development, 2017. 144(13): p. 2364–2374.

27. White, J.G., et al., The structure of the nervous system of the nematode Caenorhabditis elegans. Philos Trans R Soc Lond B Biol Sci, 1986. 314(1165): p. 1–340.

28. Chalfie, M. and J. Sulston, Developmental genetics of the mechanosensory neurons of Caenorhabditis elegans. Dev Biol, 1981. 82(2): p. 358–70.

29. Toth, M.L., et al., Neurite sprouting and synapse deterioration in the aging Caenorhabditis elegans nervous system. J Neurosci, 2012. 32(26): p. 8778–90.

30. Demontis, G.C., et al., Human Pathophysiological Adaptations to the Space Environment. Front Physiol, 2017. 8: p. 547.

31. Kaushik, S. and A.M. Cuervo, Proteostasis and aging. Nat Med, 2015. 21(12): p. 1406–15.

32. Chalfie, M., et al., Green fluorescent protein as a marker for gene expression. Science, 1994. 263(5148): p. 802–5.

33. Li, Y., et al., The lysosomal membrane protein SCAV-3 maintains lysosome integrity and adult longevity. J Cell Biol, 2016. 215(2): p. 167–185.

34. Liu, Y., et al., Autophagy-dependent ribosomal RNA degradation is essential for maintaining nucleotide homeostasis during C. elegans development. Elife, 2018. 7.

35. Wang, Z., et al., The RBG-1-RBG-2 complex modulates autophagy activity by regulating lysosomal biogenesis and function in C. elegans. J Cell Sci, 2019. 132(19).

36. Honda, Y., et al., Spaceflight and ageing: reflecting on Caenorhabditis elegans in space. Gerontology, 2014. 60(2): p. 138–42.

37. Garrett-Bakelman, F.E., et al., The NASA Twins Study: A multidimensional analysis of a year-long human spaceflight. Science, 2019. 364(6436).

38. Salzberg, Y., et al., Skin-derived cues control arborization of sensory dendrites in Caenorhabditis elegans. Cell, 2013. 155(2): p. 308–20.

39. Liang, X., et al., Sarcomeres Pattern Proprioceptive Sensory Dendritic Endings through UNC-52/Perlecan in C. elegans. Dev Cell, 2015. 33(4): p. 388–400.

40. Diaz-Balzac, C.A., et al., Muscle- and Skin-Derived Cues Jointly Orchestrate Patterning of Somatosensory Dendrites. Curr Biol, 2016. 26(17): p. 2397.

41. Zhu, T., et al., Dynein and EFF-1 control dendrite morphology by regulating the localization pattern of SAX-7 in epidermal cells. J Cell Sci, 2017. 130(23): p. 4063–4071.

42. Yang, W.K. and C.T. Chien, Beyond being innervated: the epidermis actively shapes sensory dendritic patterning. Open Biol, 2019. 9(3): p. 180257.

43. Gioran, A., P. Nicotera, and D. Bano, Impaired mitochondrial respiration promotes dendritic branching via the AMPK signaling pathway. Cell Death Dis, 2014. 5: p. e1175.

44. Xie, Y., et al., The GTP-binding protein Septin 7 is critical for dendrite branching and dendritic-spine morphology. Curr Biol, 2007. 17(20): p. 1746–51.

45. Santucci, D., et al., Evaluation of gene, protein and neurotrophin expression in the brain of mice exposed to space environment for 91 days. PLoS One, 2012. 7(7): p. e40112.

46. Willis, C.R.G., et al., Comparative Transcriptomics Identifies Altered Neuronal and Metabolic Function as Common Adaptations to Microgravity and Hypergravity in Caenorhabditis elegans. Available at SSRN: https://ssrn.com/abstract=3677482 or http://dx.doi.org/10.2139/ssrn.3677482.

47. Lesch, B.J., et al., Transcriptional regulation and stabilization of left-right neuronal identity in C. elegans. Genes Dev, 2009. 23(3): p. 345–58.

48. Jessen, N.A., et al., The Glymphatic System: A Beginner’s Guide. Neurochem Res, 2015. 40(12): p. 2583–99.

49. Weber, B. and L.F. Barros, The Astrocyte: Powerhouse and Recycling Center. Cold Spring Harb Perspect Biol, 2015. 7(12).

50. Lim, J. and Z. Yue, Neuronal aggregates: formation, clearance, and spreading. Dev Cell, 2015. 32(4): p. 491–501.

51. Ries, M. and M. Sastre, Mechanisms of Abeta Clearance and Degradation by Glial Cells. Front Aging Neurosci, 2016. 8: p. 160.

52. Ryu, H.W., et al., Simulated microgravity contributes to autophagy induction by regulating AMP-activated protein kinase. DNA Cell Biol, 2014. 33(3): p. 128–35.

53. Sambandam, Y., et al., Microgravity control of autophagy modulates osteoclastogenesis. Bone, 2014. 61: p. 125–31.

54. Blaber, E.A., M.J. Pecaut, and K.R. Jonscher, Spaceflight Activates Autophagy Programs and the Proteasome in Mouse Liver. Int J Mol Sci, 2017. 18(10).

55. Locatelli, L., et al., Mitophagy contributes to endothelial adaptation to simulated microgravity. FASEB J, 2020. 34(1): p. 1833–1845.

56. Ferranti, F., et al., Cytoskeleton modifications and autophagy induction in TCam-2 seminoma cells exposed to simulated microgravity. Biomed Res Int, 2014. 2014: p. 904396.

57. Jeong, A.J., et al., Microgravity induces autophagy via mitochondrial dysfunction in human Hodgkin’s lymphoma cells. Sci Rep, 2018. 8(1): p. 14646.

58. Fukazawa, T., et al., Simulated microgravity enhances CDDP-induced apoptosis signal via p53-independent mechanisms in cancer cells. PLoS One, 2019. 14(7): p. e0219363.

59. Clark, S.G. and C. Chiu, C. elegans ZAG-1, a Zn-finger-homeodomain protein, regulates axonal development and neuronal differentiation. Development, 2003. 130(16): p. 3781–94.

60. Gaffney, C.J., et al., Methods to assess subcellular compartments of muscle in C. elegans. J Vis Exp, 2014(93): p. e52043.

61. Djeddi, A., et al., Sperm-inherited organelle clearance in C. elegans relies on LC3-dependent autophagosome targeting to the pericentrosomal area. Development, 2015. 142(9): p. 1705–16.

62. Hadwiger, G., et al., A monoclonal antibody toolkit for C. elegans. PLoS One, 2010. 5(4): p. e10161.

